# DeepGOPlus: Improved protein function prediction from sequence

**DOI:** 10.1101/615260

**Authors:** Maxat Kulmanov, Robert Hoehndorf

**Affiliations:** Computational Bioscience Research Center, Computer, Electrical and Mathematical Sciences & Engineering Division, King Abdullah University of Science and Technology, 4700 King Abdullah University of Science and Technology, Thuwal 23955-6900, Kingdom of Saudi Arabia

## Abstract

Protein function prediction is one of the major tasks of bioinformatics that can help in wide range of biological problems such as understanding disease mechanisms or finding drug targets. Many methods are available for predicting protein functions from sequence based features, protein–protein interaction networks, protein structure or literature. However, other than sequence, most of the features are difficult to obtain or not available for many proteins thereby limiting their scope. Furthermore, the performance of sequence-based function prediction methods is often lower than methods that incorporate multiple features and predicting protein functions may require a lot of time.

We developed a novel method for predicting protein functions from sequence alone which combines deep convolutional neural network (CNN) model with sequence similarity based predictions. Our CNN model scans the sequence for motifs which are predictive for protein functions and combines this with functions of similar proteins. We evaluate the performance of DeepGOPlus on the CAFA3 dataset and significantly improve the performance of predictions of biological processes and cellular components with *F*_*max*_ of 0.47 and 0.70, respectively, using only the amino acid sequence of proteins as input. DeepGOPlus can annotate around 40 protein sequences per second, thereby making fast and accurate function predictions available for a wide range of proteins.

## INTRODUCTION

Prediction of protein functions is a major task in bioinformatics that is important in understanding the role of proteins in disease pathobiology, the functions of metagenomes, or finding drug targets. A wide range of methods have been developed for predicting protein functions computationally (Kahanda and Ben-Hur, 2017, Kulmanov *et al.*, 2017, Radivojac *et al.*, 2013, You *et al.*, 2018a, b). Protein functions can be predicted from protein sequences (Kulmanov *et al.*, 2017, Radivojac *et al.*, 2013, You *et al.*, 2018a, b), protein–protein interactions (PPI) (Kulmanov *et al.*, 2017), protein structures (Yang *et al.*, 2014), biomedical literature, and other features (Kahanda and Ben-Hur, 2017, You *et al.*, 2018a). Sequence-based methods employ sequence similarity, search for sequence domains, or multi-sequence alignments to infer functions. As proteins rarely function on their own, protein–protein interactions can be a good predictor for complex biological processes to which proteins contribute. Although it is experimentally challenging to identify protein structures, they are crucial in understanding what proteins are capable of doing. Literature may contribute to function predicting because it may contain explicit descriptions of protein functions or describe properties of proteins that are predictive of protein functions indirectly. Overall, many of these features are available only for a small number of proteins, while a protein’s amino acid sequence can be identified for most proteins. Therefore, methods that accurately predict protein functions from sequence alone may be the most general and applicable to proteins that have not been extensively studied.

Proteins with similar sequence tend to have similar functions (Radivojac *et al.*, 2013). Therefore, a basic way of predicting functions for new sequences is to find the most similar sequences with known functional annotations and transfer their annotations. Another approach is to search for specific sequence motifs which are associated with some function; for example, InterProScan (Mitchell *et al.*, 2014) is a tool which can help to find protein domains and families. The domains and families can the be used to infer protein functions.

Recent developments in deep feature learning methods brought many methods which can learn protein sequence features. In 2017, we developed DeepGO (Kulmanov *et al.*, 2017) as one of first deep learning models which can predict protein functions using the protein amino acid sequence and interaction networks. Since 2017, many successor methods became available that achieve better predictive performance (You *et al.*, 2018a, b).

DeepGO suffers from several limitations. First, it can only predict functions for proteins with a sequence length less than 1002 and which do not contain “ambiguous” amino acids such as unions or unknowns. While around 90% of protein sequences in UniProt satisfy these criteria, it also means that DeepGO could not predict functions for about 10% of proteins. Second, due to computational limitations, DeepGO can only predict around 2,000 functions out of more than 45,000 which are currently in the Gene Ontology (GO) (Ashburner *et al.*, 2000). Third, DeepGO uses interaction network features which are not available for all proteins. Specifically, for novel or uncharacterized proteins, only the sequence may be known and not any additional information such as the protein’s interactions or mentions in literature. Finally, DeepGO was trained and evaluated on randomly drawn training, validation, and testing sets. However, such models may overfit to particular features in the training data and may not yield adequate results in real prediction scenarios. Consequently, challenges such as the Critical Assessment of Function Annotation (CAFA) (Radivojac *et al.*, 2013) use a time-based evaluation where training and predictions are fixed and evaluated after some time has elapsed on predictions that became available in that time. DeepGO did not achieve the same performance in the CAFA3 challenge as it had in our own experiments.

Here, we extend and improve DeepGO overcoming its main limitations related to sequence length, missing features, and number of predicted classes. We increased the model’s input length to 2,000 amino acids and now cover more than 99% of sequences in UniProt. Furthermore, our new model’s architecture allows us to split longer sequences and scan smaller chunks to predict functions. We also remove features derived from interaction networks because only a small number of proteins have such network information. Instead, we combine our neural network predictions with methods based on sequence similarity to capture orthology and, indirectly, some interaction information. Through this step we also overcome the limitation in the number of classes to predict and we can, in theory, predict any GO class that has ever been used in an experimental annotation. To avoid overfitting of our model, we substantially decreased our model’s capacity by replacing the amino acid trigram embedding layer with a one-hot encoding and removing our hierarchical classification layer. Moreover, by using a single model with less parameters we significantly improved the runtime of the model. In average, DeepGOPlus can annotate 40 proteins per second on ordinary hardware. Overall, with these improvements, our model can now perform *de novo* predictions for any protein with available sequence.

In our evaluation we exactly reproduce the CAFA3 evaluation by training our model using only data provided by CAFA3 as training data and evaluating on the CAFA3 testing data. We compare DeepGOPlus with our baseline methods including DeepGO and two best-performing protein function prediction methods, GOLabeler (You *et al.*, 2018b) and DeepText2GO (You *et al.*, 2018a). GOLabeler mainly uses sequence-based features, DeepGO uses interaction network features, and DeepText2GO uses features extracted from literature in addition to sequence-based ones. In terms of *F*_*max*_ measure, we outperform all methods in predicting biological processes and cellular components. Notably, our model significantly improves predictions of biological process annotations with an *F*_*max*_ of 0.474.

## MATERIALS AND METHODS

### Datasets and Gene Ontology

We use two datasets to evaluate our approach. Firstly, we downloaded CAFA3 challenge (Radivojac *et al.*, 2013) training sequences and experimental annotations published on September, 2016 and test benchmark published on 15th November 2017 which was used to evaluate protein function prediction methods submitted to the challenge. According to CAFA3, the annotations with evidence codes: EXP, IDA, IPI, IMP, IGI, IEP, TAS, or IC are considered to be experimental. The training set includes all proteins with experimental annotations known before September, 2016 and the test benchmark contains no-knowledge proteins which gained experimental annotation between September, 2016 and November 2017. Similar time based splits were used in all previous CAFA challenges.

We propagate annotations using the hierarchical structure of the Gene Ontology (GO) (Ashburner *et al.*, 2000). We use the version of GO released on 1 June 2016. The version has 10,693 molecular function (MFO), classes, 29,264 biological process (BPO) classes and 4,034 cellular component (CCO) classes. This version is also used to evaluate CAFA3 predictions. While propagating annotations, we consider all types of relations between classes. For instance, if a protein *P* is annotated with a class *C* which has a part-of relation to a class *D*, then we annotate *P* with the class *D*. This procedure is repeated until no further annotation can be propagated. After this, we count the number of annotated proteins for each GO class and select all classes with 50 or more annotations for our prediction model. The statistics with the number of classes in Table 1 represent how many classes we can predict using our deep neural network model.

**Table 1.** The number of protein sequences with experimental annotations in CAFA3 and 2016 datasets grouped by sub-ontologies.

| Dataset | Statistic | MFO | BPO | CCO | All |
| --- | --- | --- | --- | --- | --- |
| CAFA3 | Training size | 36,110 | 53,500 | 50,596 | 66,841 |
| CAFA3 | Testing size | 1,137 | 2,392 | 1,265 | 3,328 |
| CAFA3 | Number of classes | 677 | 3992 | 551 | 5,220 |
| 2016 | Training size | 34,488 | 51,716 | 49,346 | 65,028 |
| 2016 | Testing size | 679 | 1,434 | 1,148 | 1,788 |
| 2016 | Number of classes | 652 | 3,904 | 545 | 5,101 |

Secondly, to compare with other methods for function prediction such as DeepText2GO (You *et al.*, 2018a) and GoLabeler (You *et al.*, 2018b) we downloaded SwissProt reviewed proteins published on January, 2016 and October, 2016. We use all experimental annotations before January 2016 as a training set and experimental annotations collected between January and October 2016 as testing set. We filter the testing set with 23 target species which are in CAFA evaluation set. Table 1 summarizes both datasets.

### Baseline comparison methods

#### Naive approach

It is possible to get comparable prediction results just by assigning the same GO classes to all proteins based on annotation frequencies. This happens due to the hierarchical structure of GO which, after the propagation process, results in many annotations at high-level classes. In CAFA, this approach is called “naive” approach and is used as one of the baseline methods to compare function predictions. Here, each query protein *p* is annotated with the GO classes with a prediction scores computed as:

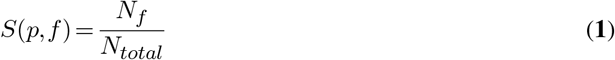

where *f* is a GO class, *N*_*f*_ is a number of training proteins annotated by GO class *f* and *N*_*total*_ is a total number of training proteins.

#### DiamondBLAST

Another baseline method is based on sequence similarity score obtained by BLAST (Altschul *et al.*, 1997). The idea is to find similar sequences from the training set and transfer an annotation from the most similar. We use the normalized bitscore as prediction score for a query sequence *q*:

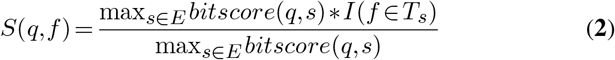

where *E* is a set of similar sequences filtered by e-value of 0.001, *T*_*s*_ is a set of true annotations of a protein with sequence *s* and *I* is an identity function which returns 1 if the condition is true and 0 otherwise.

#### DiamondScore

The DiamondScore is very similar to the DiamondBLAST approach. The only difference is that we normalize the sum of the bitscores of similar sequences. We compute prediction scores using the formula:

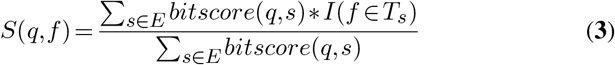

#### DeepGO

DeepGO (Kulmanov *et al.*, 2017) was developed by us previously and it is one of the first methods which learns sequence features with a deep learning model and combines it with PPI network features to predict protein functions. It also uses a hierarchical classifier to output predictions consistent with structure of GO. Here we trained three separate models for three parts of GO mainly because of the computational costs involved in training larger models. We use our previously reported optimal parameters and set of functions to train new models with our current datasets. With DeepGO, we trained and predicted 932 BPO, 589 MFO and 436 CCO classes.

#### GOLabeler and DeepText2GO

Currently the best performing methods for function prediction task are GOLabeler (You *et al.*, 2018b) and DeepText2GO (You *et al.*, 2018a), both developed by the same group. GOLabeler achieved some of the best results in the preliminary evaluation for all three subontologies of GO in the CAFA3 challenge. It is an ensemble method which combines several approaches and predicts functions mainly from sequence features. DeepText2GO improves the results achieved by GOLabeler by extending their ensemble with models that predict functions from literature.

Our second dataset is specifically designed to compare our results with these two methods. Since we use same training and testing data, we directly compare our results with the results reported in their papers.

### Model Training

We use Tensorflow (Abadi *et al.*, 2016) to build and train our neural network model. Our model was trained on Nvidia Titan X and P6000 GPUs with 12-24Gb of RAM.

### Evaluation

To evaluate our predictions we use the CAFA (Radivojac *et al.*, 2013) evaluation metrics *F*_*max*_ and *S*_*min*_ (Radivojac and Clark, 2013). In addition, we report area under the precision-recall curve (AUPR) which is a reasonable measure for evaluating predictions with high class imbalance (Davis and Goadrich, 2006).

*F*_*max*_ is a maximum protein-centric F-measure computed over all prediction thresholds. First, we compute average precision and recall using the following formulas:

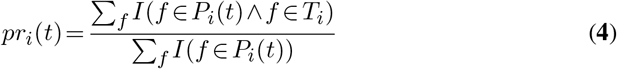

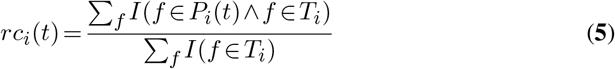

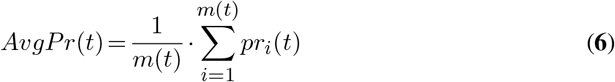

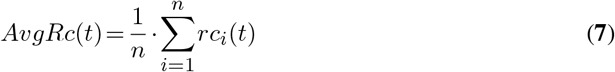

where *f* is a GO class, *T*_*i*_ is a set of true annotations, *P*_*i*_(*t*) is a set of predicted annotations for a protein *i* and threshold *t*, *m*(*t*) is a number of proteins for which we predict at least one class, *n* is a total number of proteins and *I* is an identity function which returns 1 if the condition is true and 0 otherwise. Then, we compute the *F*_*max*_ for all possible thresholds:

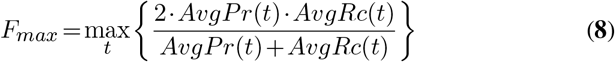

*S*_*min*_ computes semantic distance between real and predicted annotations based on information content of the classes. The information content *IC*(*c*) is computed based on the annotation probability of the class *c*:

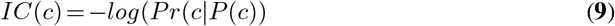

where *P* (*c*) is a set of parent classes of the class *c*. The *S*_*min*_ is computed using the following formulas:

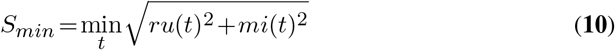

where *ru*(*t*) is average remaining uncertainty and *mi*(*t*) is average misinformation:

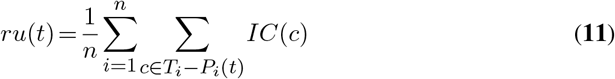

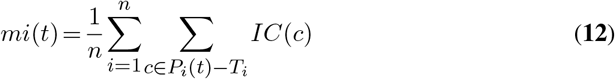

## RESULTS

### DeepGOPlus Learning Model

In DeepGOPlus, we combine sequence similarity and sequence motifs in a single predictive model. To learn sequence motifs that are predictive of protein functions, we use one-dimensional convolutional neural networks (CNNs) over protein amino acid sequence to learn sequence patterns or motifs. Figure 1 describes the architecture of our deep learning model. First, the input sequence is converted to a one-hot encoded representation of size 21 × 2000, where a one-hot vector of length 21 represents an amino acid (AA) and 2,000 is the input length. Sequences with a length less than 2,000 are padded with zeros and longer sequences are split into smaller chunks with less than 2,000 AAs. This input is passed to a set of CNN layers with different filter sizes of 8,16,…,128. Each of the CNN layers has 512 filters which learn specific sequence motifs of a particular size. Each filter is scanning the sequence and their maximum score is pooled using a MaxPooling layer. In total, we generate a feature vector of size 8,192 where each value represents a score that indicates the presence of a relevant sequence motif. This vector is passed to the fully connected classification layer which outputs the predictions. To select the best parameters and hyperparameters for our deep learning model, we extensively searched for optimal combinations of parameters such as filter sizes, number of filters and depth of dense layers based on a validation set loss. We report the list of parameters and validation losses in Supplementary Table 1.

**Figure 1.**
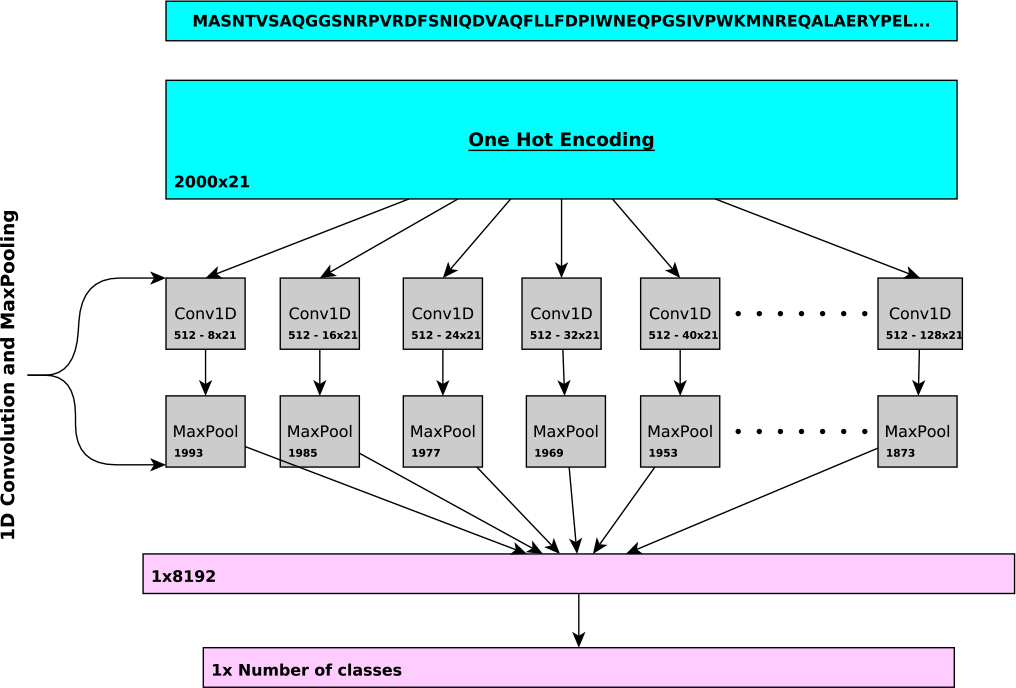
Overview of the CNN in DeepGOPlus. The CNN uses multiple filters of variable size to detect the presence of sequence motifs in the input amino acid sequence.

DeepGOPlus combines the neural network model predictions with predictions based on sequence similarity. First, we find similar sequences from a training set using Diamond (Buchfink *et al.*, 2014) with an *e*-value of 0.001 and obtain a bitscore for every similar sequence. We transfer all annotations of similar sequences to a query sequence with prediction scores computed using the bitscores. For a set of similar sequences *E* of the query sequence *q*, we compute the prediction score for a GO class *f* as

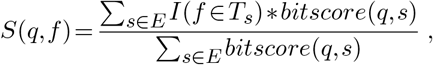

where *T*_*s*_ is a set of true annotations of the protein with sequence *s*. Then, to compute the final prediction scores of DeepGOPlus, we combine the two prediction scores using a weighted sum model (Fishburn, 1967):

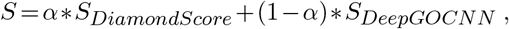

where 0≤*α*≤1 is a weight parameter which balances the relative importance of the two prediction methods.

### Evaluation and comparison

We evaluate DeepGOPlus using two datasets. First, we use the latest CAFA3 (Radivojac *et al.*, 2013) challenge dataset and compare our method with baseline methods such as Naive predictions, BLAST, and our previous deep learning model DeepGO. We use two strategies for predicting functions based on sequence similarity computed with the Diamond tool (Buchfink *et al.*, 2014) (which is a faster implementation of the BLAST algorithm). We call them DiamondBLAST and DiamondScore. DiamondBLAST considers only the most similar sequence whereas DiamondScore predicts functions using all similar sequences returned by Diamond. We also report the performance of using only our neural network model (labeled as DeepGOCNN). We find that with the DiamondScore approach, we can outperform DeepGO predictions in MFO and achieve comparable results in BPO and CCO evaluations while DeepGOCNN gives better predictions in CCO. We achieve the best performance in all three subontologies with our DeepGOPlus model which combines the DiamondScore and DeepGOCNN. Table 2 summarizes the performance of the models.

**Table 2.** The comparison of performance on the first CAFA3 challenge dataset.

| Method | $F_{max}$ | | | $S_{min}$ | | | AUPR | | |
| --- | --- | --- | --- | --- | --- | --- | --- | --- | --- |
|  | MFO | BPO | CCO | MFO | BPO | CCO | MFO | BPO | CCO |
| Naive | 0.290 | 0.357 | 0.562 | 10.733 | 25.028 | 8.465 | 0.130 | 0.254 | 0.456 |
| DiamondBLAST | 0.431 | 0.399 | 0.506 | 10.233 | 25.320 | 8.800 | 0.178 | 0.116 | 0.142 |
| DiamondScore | 0.509 | 0.427 | 0.557 | 9.031 | 22.860 | 8.198 | 0.340 | 0.267 | 0.335 |
| DeepGO | 0.393 | 0.435 | 0.565 | 9.635 | 24.181 | 9.199 | 0.303 | 0.385 | 0.579 |
| DeepGOCNN | 0.420 | 0.378 | 0.607 | 9.711 | 24.234 | 8.153 | 0.355 | 0.323 | 0.616 |
| DeepGOPlus | **0.547** | **0.470** | **0.625** | **8.660** | **22.514** | **7.804** | **0.489** | **0.405** | **0.631** |

**Table 3.**
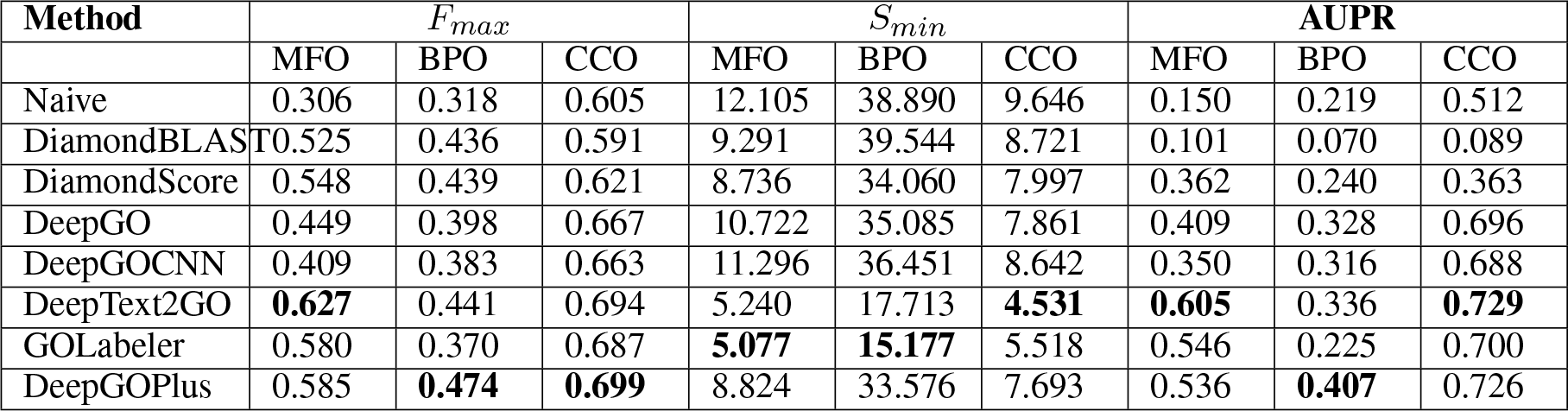
The comparison of performance on the second dataset generated by a time-based split.

| Method | $F_{max}$ | | | $S_{min}$ | | | AUPR | | |
| --- | --- | --- | --- | --- | --- | --- | --- | --- | --- |
|  | MFO | BPO | CCO | MFO | BPO | CCO | MFO | BPO | CCO |
| Naive | 0.306 | 0.318 | 0.605 | 12.105 | 38.890 | 9.646 | 0.150 | 0.219 | 0.512 |
| DiamondBLAST | 0.525 | 0.436 | 0.591 | 9.291 | 39.544 | 8.721 | 0.101 | 0.070 | 0.089 |
| DiamondScore | 0.548 | 0.439 | 0.621 | 8.736 | 34.060 | 7.997 | 0.362 | 0.240 | 0.363 |
| DeepGO | 0.449 | 0.398 | 0.667 | 10.722 | 35.085 | 7.861 | 0.409 | 0.328 | 0.696 |
| DeepGOCNN | 0.409 | 0.383 | 0.663 | 11.296 | 36.451 | 8.642 | 0.350 | 0.316 | 0.688 |
| DeepText2GO | **0.627** | 0.441 | 0.694 | 5.240 | 17.713 | **4.531** | **0.605** | 0.336 | **0.729** |
| GOLabeler | 0.580 | 0.370 | 0.687 | **5.077** | **15.177** | 5.518 | 0.546 | 0.225 | 0.700 |
| DeepGOPlus | 0.585 | **0.474** | **0.699** | 8.824 | 33.576 | 7.693 | 0.536 | **0.407** | 0.726 |

To compare our approach with the state of the art methods GOLabeler (You *et al.*, 2018b) and DeepText2GO (You *et al.*, 2018a), we generate a second dataset which uses data obtained at the same dates as the other methods so that we can generate a time-based split of training and testing data. Both methods train on experimental function annotations that appeared before January 2016 and test on annotations which were asserted between January 2016 and October 2016.

Furthermore, we use the same version of GO and follow the CAFA3 challenge procedures to process the data. As a result, we can directly compare our evaluation results with the other methods. In this evaluation, DeepGOPlus gives the best results for BPO and CCO in terms of *F*_*max*_ measure and ranks second in the MFO evaluation (after DeepText2GO). However, it is important to note that DeepText2GO uses features extracted from literature in addition to sequence based features while DeepGOPlus predictions are only based on protein sequence. Notably, our method significantly increased performance of predictions of BPO classes in both evaluation datasets.

Due to large number of available sequences, analyzing sequences require both accurate and fast prediction methods. Specifically, function prediction is a crucial step in interpretation of newly-sequenced genomes or meta-genomes. While we have compared DeepGOPlus in terms of prediction performance, we could not compare the running time of the models because the runtime of prediction models is rarely reported. With DeepGOPlus, 40 protein sequences can be annotated per second using a single Intel(R) Xeon(R) E5-2680 CPU and Nvidia P6000 GPU.

### Implementation and availability

DeepGOPlus is available as free software at https://github.com/bio-ontology-research-group/deepgoplus. We also publish training and testing data used to generate evaluation and results at http://deepgoplus.bio2vec.net/data/. Furthermore, DeepGOPlus is available through a REST API at http://deepgoplus.bio2vec.net.

## DISCUSSION

DeepGOPlus is a fast and accurate tool to predict protein functions from protein sequence alone. Our model overcomes several limitations of other methods and our own DeepGO model (Kulmanov *et al.*, 2017). In particular, DeepGOPlus has no limits on the length of the amino acid sequence and can therefore be used for the genome-scale annotation of protein functions, in particular in newly-sequenced organisms. DeepGOPlus also makes no assumptions on the taxa or kingdom to which a protein belongs, therefore enabling, for example, function prediction for meta-genomics in which proteins from different kingdoms may be mixed. Furthermore, DeepGOPlus is fast and can annotate several thousand proteins in minutes even on single CPUs, further enabling its application in metagenomics or for projects in which a very large number of proteins with unknown functions are identified. While we initially expected the absence of features derived from interaction networks to impact predictive performance, we found that we can achieve even higher prediction accuracy with our current model; additionally, our model is not limited by unbalanced or missing information about protein-protein interactions.

In DeepGOPlus, we combine similarity-based search to proteins with known functions and motif-based function prediction, and this combination gives us overall the best predictive performance. However, DeepGOPlus can also be applied using only sequence motifs; in particular when annotating novel proteins for which no similar proteins with known functions exist, our motif-based model would be most suitable.

In the future, we plan to incorporate additional features. While related methods use features that can be derived only for known proteins, such as information obtained from literature or interaction networks, DeepGOPlus will rely primarily on features that can be derived from amino acid sequences to ensure that the model can be applied as widely as possible. Possible additional information that may improve DeepGOPlus in the future is information about protein structure, in particular as structure prediction methods are improving significantly (Wang *et al.*, 2017).

