## Supplementary Table 1 for "DeepGOPlus: Improved protein function prediction from sequence"

### Supplementary materials: DeepGOPlus: Improved protein function prediction from sequence

April 18, 2019

| # | MaxKernel | Hidden Layers | Filters | Valid. Loss | Test Loss |
|---|---|---|---|---|---|
| 1 | 32 | 1 | 32 | 0.04085 | 0.024466 |
| 2 | 64 | 1 | 32 | 0.04032 | 0.024393 |
| 3 | 128 | 1 | 32 | 0.04038 | 0.024314 |
| 4 | 256 | 1 | 32 | 0.04008 | 0.024610 |
| 5 | 512 | 1 | 32 | 0.04019 | 0.025172 |
| 6 | 32 | 2 | 32 | 0.04187 | 0.024399 |
| 7 | 64 | 2 | 32 | 0.04194 | 0.024147 |
| 8 | 128 | 2 | 32 | 0.04125 | 0.026296 |
| 9 | 256 | 2 | 32 | 0.04054 | 0.025792 |
| 10 | 512 | 2 | 32 | 0.04134 | 0.024686 |
| 11 | 32 | 3 | 32 | 0.04173 | 0.024688 |
| 12 | 64 | 3 | 32 | 0.04215 | 0.025352 |
| 13 | 128 | 3 | 32 | 0.04110 | 0.025408 |
| 14 | 256 | 3 | 32 | 0.04109 | 0.024851 |
| 15 | 512 | 3 | 32 | 0.04104 | 0.026999 |
| 16 | 32 | 1 | 64 | 0.04028 | 0.023811 |
| 17 | 64 | 1 | 64 | 0.04016 | 0.024081 |
| 18 | 128 | 1 | 64 | 0.04024 | 0.024422 |
| 19 | 256 | 1 | 64 | 0.03986 | 0.025013 |
| 20 | 512 | 1 | 64 | 0.04039 | 0.024882 |
| 21 | 32 | 2 | 64 | 0.04169 | 0.024334 |
| 22 | 64 | 2 | 64 | 0.04117 | 0.025407 |
| 23 | 128 | 2 | 64 | 0.04080 | 0.026773 |
| 24 | 256 | 2 | 64 | 0.04043 | 0.025632 |
| 25 | 512 | 2 | 64 | 0.04039 | 0.026199 |
| 26 | 32 | 3 | 64 | 0.04160 | 0.024624 |
| 27 | 64 | 3 | 64 | 0.04142 | 0.026038 |
| 28 | 128 | 3 | 64 | 0.04029 | 0.025585 |
| 29 | 256 | 3 | 64 | 0.04016 | 0.025938 |

| | | | | | |
|---|---|---|---|---|---|
| 30 | 512 | 3 | 64 | 0.04052 | 0.026569 |
| 31 | 32 | 1 | 128 | 0.03996 | 0.023886 |
| 32 | 64 | 1 | 128 | 0.03999 | 0.024146 |
| 33 | 128 | 1 | 128 | 0.03990 | 0.024231 |
| 34 | 256 | 1 | 128 | 0.03945 | 0.025394 |
| 35 | 512 | 1 | 128 | 0.04002 | 0.024113 |
| 36 | 32 | 2 | 128 | 0.04131 | 0.024404 |
| 37 | 64 | 2 | 128 | 0.04116 | 0.024885 |
| 38 | 128 | 2 | 128 | 0.04000 | 0.026053 |
| 39 | 256 | 2 | 128 | 0.04000 | 0.026025 |
| 40 | 512 | 2 | 128 | 0.04037 | 0.023884 |
| 41 | 32 | 3 | 128 | 0.04100 | 0.025189 |
| 42 | 64 | 3 | 128 | 0.04031 | 0.025051 |
| 43 | 128 | 3 | 128 | 0.03995 | 0.025764 |
| 44 | 256 | 3 | 128 | 0.04135 | 0.024185 |
| 45 | 512 | 3 | 128 | 0.04008 | 0.024312 |
| 46 | 32 | 1 | 256 | 0.03942 | 0.023801 |
| 47 | 64 | 1 | 256 | 0.03933 | 0.024252 |
| 48 | 128 | 1 | 256 | 0.03897 | 0.024517 |
| 49 | 256 | 1 | 256 | 0.03950 | 0.024034 |
| 50 | 512 | 1 | 256 | 0.04146 | 0.024329 |
| 51 | 32 | 2 | 256 | 0.04050 | 0.024755 |
| 52 | 64 | 2 | 256 | 0.04026 | 0.025917 |
| 53 | 128 | 2 | 256 | 0.03992 | 0.025778 |
| 54 | 256 | 2 | 256 | 0.03995 | 0.025696 |
| 55 | 512 | 2 | 256 | 0.04071 | 0.025106 |
| 56 | 32 | 3 | 256 | 0.04129 | 0.025048 |
| 57 | 64 | 3 | 256 | 0.03997 | 0.024811 |
| 58 | 128 | 3 | 256 | 0.03918 | 0.025124 |
| 59 | 256 | 3 | 256 | 0.03920 | 0.026214 |
| 60 | 512 | 3 | 256 | 0.04045 | 0.026019 |
| 61 | 32 | 1 | 512 | 0.03912 | 0.024001 |
| 62 | 64 | 1 | 512 | 0.03881 | 0.024483 |
| 63 | 128 | 1 | 512 | **0.03880** | 0.024031 |
| 64 | 256 | 1 | 512 | 0.03966 | 0.024428 |
| 65 | 512 | 1 | 512 | 0.04201 | 0.024437 |
| 66 | 32 | 2 | 512 | 0.03929 | 0.025065 |
| 67 | 64 | 2 | 512 | 0.03913 | 0.025310 |
| 68 | 128 | 2 | 512 | 0.03945 | 0.024313 |
| 69 | 256 | 2 | 512 | 0.03954 | 0.024128 |
| 70 | 512 | 2 | 512 | 0.04030 | 0.024147 |
| 71 | 32 | 3 | 512 | 0.03947 | **0.023562** |
| 72 | 64 | 3 | 512 | 0.03925 | 0.025331 |
| 73 | 128 | 3 | 512 | 0.03902 | 0.024775 |
| 74 | 256 | 3 | 512 | 0.03969 | 0.024065 |
| 75 | 512 | 3 | 512 | 0.03971 | 0.025466 |

Table 1: Different parameters used to tune the model
